# Web-based Tool Validation for Antimicrobial Resistance Prediction: An Empirical Comparative Analysis

**DOI:** 10.1101/2022.12.08.519699

**Authors:** Sweta Padma Routray, Swayamprabha Sahoo, Debasish Swapnesh Kumar Nayak, Sejal Shah, Tripti Swarnkar

**Affiliations:** Center for Biotechnology, Siksha ‘O’ Anusandhan Deemed to be University, Bhubaneswar, India-751003, Email.Id; Department of Computer Science & Engineering, Institute of Technical Education and Research, Siksha ‘O’ Anusandhan Deemed to be University, Bhubaneswar, India-751030, Email.Id; Department of Computer Science and Bioscience, Faculty of Technology, Marwadi University, Rajkot, Gujarat, India-360003, Email.Id; Department of Computer Application, Institute of Technical Education and Research, Siksha ‘O’ Anusandhan Deemed to be University, Bhubaneswar, India-751030, Email.Id

**Keywords:** Antimicrobial resistance (AMR), Antibiotic Resistance Genes (ARGs), ResFinder, KmerResistance, ResfinderFG

## Abstract

Global public health is seriously threatened by Antimicrobial Resistance (AMR), and there is an urgent need for quick and precise AMR diagnostic tools. The prevalence of novel Antibiotic Resistance Genes (ARGs) has increased substantially during the last decade, owing to the recent burden of microbial sequencing. The major problem is extracting vital information from the massive amounts of generated data. Even though there are many tools available to predict AMR, very few of them are accurate and can keep up with the unstoppable growth of data in the present. Here, we briefly examine a variety of AMR prediction tools that are available. We highlighted three potential tools from the perspective of the user experience that is preferable web-based AMR prediction analysis, as a web-based tool offers users accessibility across devices, device customization, system integration, eliminating the maintenance hassles, and provides enhanced flexibility and scalability. By using the *Pseudomonas aeruginosa* Complete Plasmid Sequence (CPS), we conducted a case study in which we identified the strengths and shortcomings of the system and empirically discussed its prediction efficacy of AMR sequences, ARGs, amount of information produced and visualisation. We discovered that ResFinder delivers a great amount of information regarding the ARGS along with improved visualisation. KmerResistance is useful for identifying resistance plasmids, obtaining information about related species and the template gene, as well as predicting ARGs. ResFinderFG does not provide any information about ARGs, but it predicts AMR determinants and has a better visualisation than KmerResistance.

**Author summary:** AMR is the capacity of microorganisms to survive or grow in the presence of drugs intended to stop them or kill them. Consequently, there is an increase in the Burden of disease, death rates, and the cost of healthcare, making it a serious global threat to both human and animal health. Next-Generation Sequencing (NGS) based molecular monitoring can be a real boon to phenotypic monitoring of AMR. Researchers face difficult challenges in terms of producing, managing, analysing, and interpreting massive amounts of sequence data. There are many tools available to predict AMR, but only a small number of them are reliable and able to keep up with the current rate of unstoppable data growth. Each tool has specific benefits and drawbacks of its own. Our research offers a comprehensive overview of the outcomes produced by three different tools, enabling users to choose the tool that best suits their requirements.

## Introduction

The world’s most severe public health concern is the rapid growth of resistant superbugs, as well as the ongoing battle between bactericidal drugs and extensively drug-resistant bacteria. Antibiotic overuse in both medical and agricultural contexts has aided in the development of multidrug-resistant (MDR) bacteria. Unfortunately, many antibiotics lack specificity, killing pathogenic and non-pathogenic bacteria indiscriminately and leading to antibiotic-associated illnesses (1). The innovation of new anti-bacterial to treat resistant infections is a major goal in healthcare, yet no obvious answer to this problem has been identified. These antibiotic-resistant genes allow bacteria to resist antibiotics in a variety of ways, including the activation of efflux pumps, antibiotic molecule degradation by enzymes, and chemical alteration (ribosome and cell wall) to protect antibiotic-targeted cellular targets. These resistance mechanisms, when combined, represent a danger to the therapeutic effectiveness of antibiotics (2,3). The World Health Organization (WHO) lists AMR as one of the top ten global public health threats to humanity (4). Drug-resistant diseases are also expected to kill ten million people every year by 2050. This indicates that drug-resistant diseases will result in more fatalities than road accidents, diabetes, and cancer combined (5). Antibiotic formulation, testing, and screening are resource-intensive and expensive, limiting possible treatment choices for resistant bacterial species.

ARGs have the ability to threaten public health (6). ARGs are commonly present on transposons or plasmids, and they can be transferred from one cell to another by transduction, transformation, or conjugation. Resistance spreads rapidly within a bacterial population and among different types of bacteria due to gene transfer. This is known as horizontal gene transmission (7). The detection of these genes is essential for identifying resistant strains, validating non-susceptible phenotypes, and better understanding resistance epidemiology (8). Phenotypic assays have traditionally been used to detect AMR. The criterion for determining antibiotic sensitivity is diffusion-based or standardised dilution in vitro Antibiotic Susceptibility Test (AST), and much research and testing has been done to link AST findings with treatment response. Resistance surveillance and in some cases clinical therapeutic guidance are increasingly using molecular approaches. These approaches include everything from PCR-based resistance element detection to mass spectrometry-based methods(9). Sequencing has become a viable approach for routine bacterial characterization due to the enhanced accessibility and cheaper cost of NGS. Over the past few years, it has significantly improved our ability to combat AMR. Although NGS-based technologies can detect practically any known AMR gene or mutation and explore new variations of known AMR determinants, they have largely replaced traditional genotypic approaches for AMR identification (10–12).

Furthermore, as new phenotypic AMR determinants are discovered, sequence data can be continuously stored and re-analyzed. The primary drawback of any genotypic AST approach is that it can detect only proven AMR mechanisms, while resistance induced by novel mechanisms and/or gene expression regulation (heteroresistance, increased efflux pump expression, etc.) cannot be detected (13).

The prediction of AMR in NGS data using a simple and quick method is essential for source tracking, infectious disease detection, diagnostics, and epidemiological surveillance, and additional research is highly required(14). The development of tools for continuously monitoring AMR globally has become more important because these tools may provide useful information to assist healthcare professionals in developing stewardship programmes and implementing public health measures. It is also required to create an infrastructure and network to support this huge amount of data (15). AMR is detected computationally by querying input DNA or amino acid sequence data for the existence of a pre-determined set of AMR determinants provided in AMR reference databases using a search algorithm (Figure 1) (16). Numerous drug resistance prediction tools have been developed and made publicly available online over the last couple of years (17). There is however, a requirement for the standardization of Tools. Our research has the ability to determine the relative advantages and drawbacks of the tools of the predictions generated by the various tools.

**Figure 1:**
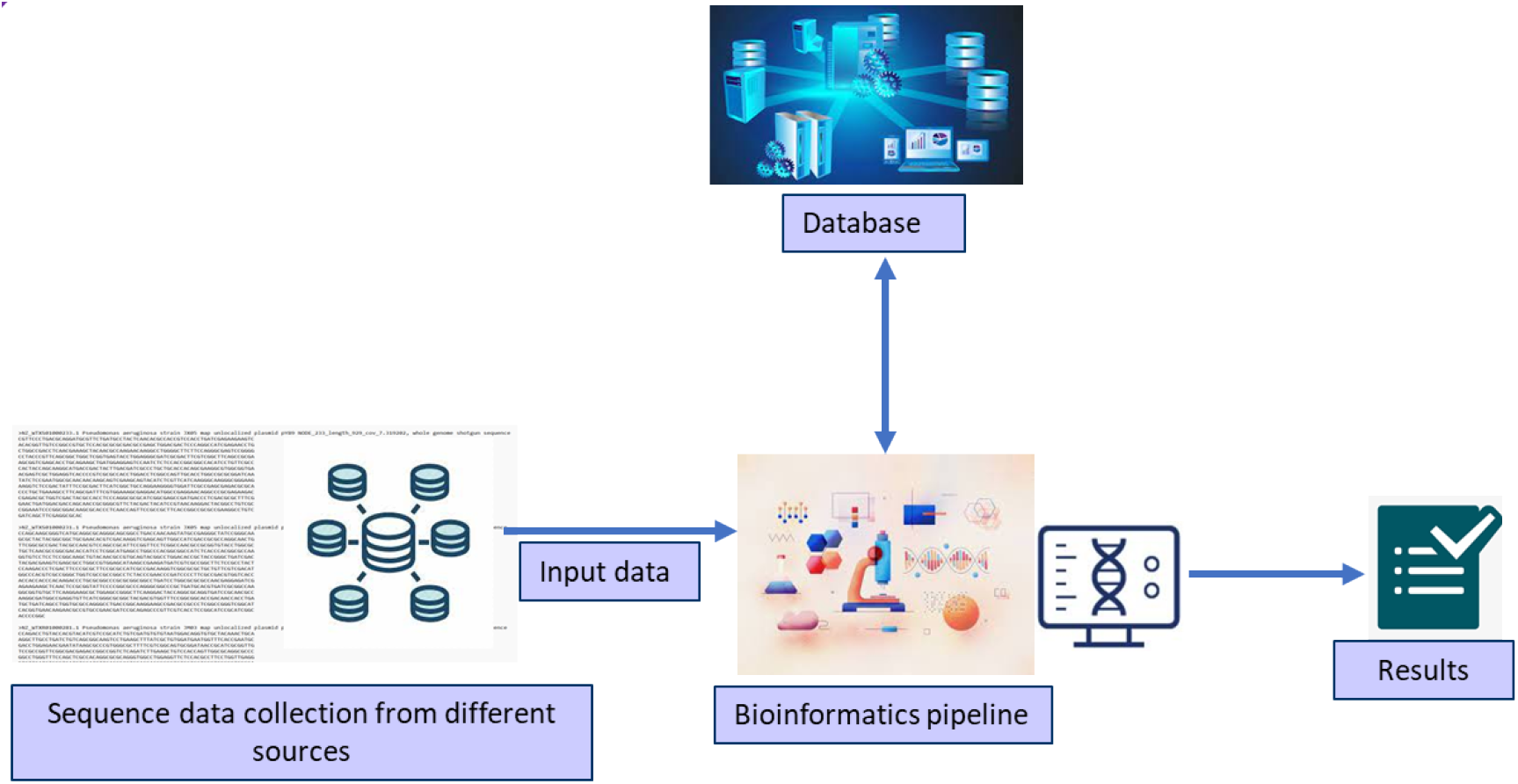
Working principle of AMR prediction tools.

## Results

This study provides better insights into the advantages and drawbacks of the tools, such as user-friendliness, to determine which tool has the best visualisation and which tool offers the maximum information.

### Results have been discussed with the following aspects

➢ Comparison based on the output resistant plasmid sequences
➢ Comparison based on the output Antibiotic Resistance Genes (ARGs)
➢ Comparison based on the amount of information provided and result visualisation
➢ In detailed result of each tool

### Comparison based on the output resistant plasmid sequences

ResFinder, KmerResistance, and ResFinderFG were each provided 25 plasmid sequences to analyse, and these three tools examined all sequences. All three tools predicted AMR for the five common sequences, whereas ResFinder and KmerResistance both predicted 13 sequences that are shared by both. ResFinderFG predicted AMR in only six sequences (Figure 4).

**Figure 2.**
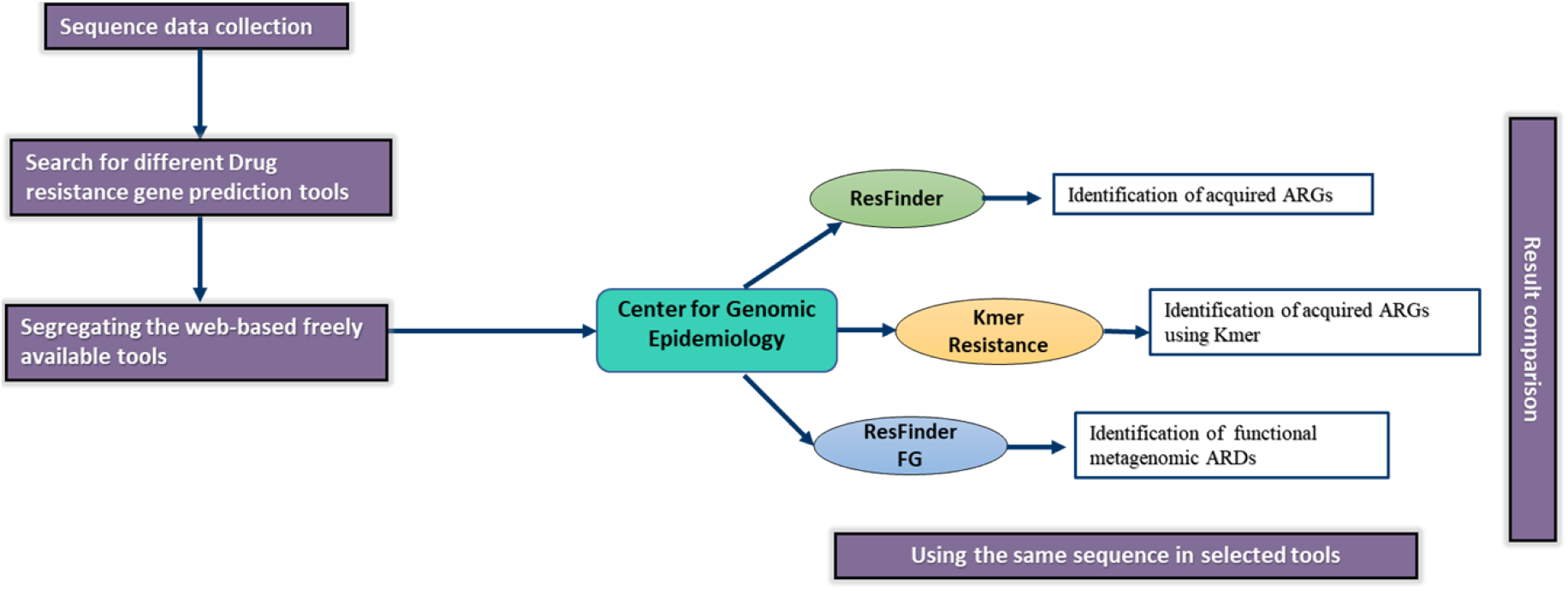
An overview of the AMR prediction tools pipeline

**Figure 3.**
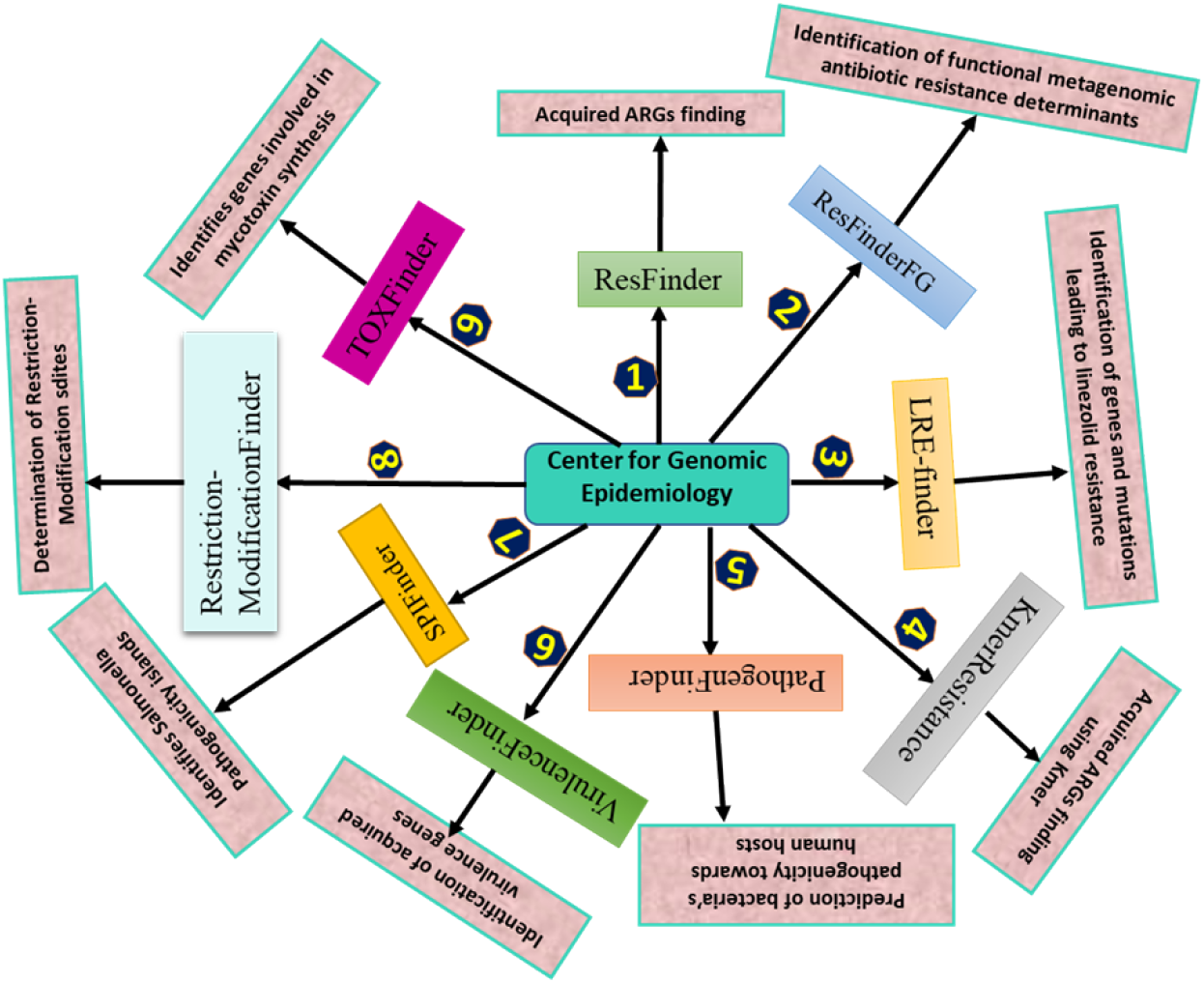
Center for Genomic Epidemiology based phenotyping tools.

**Figure 4.**
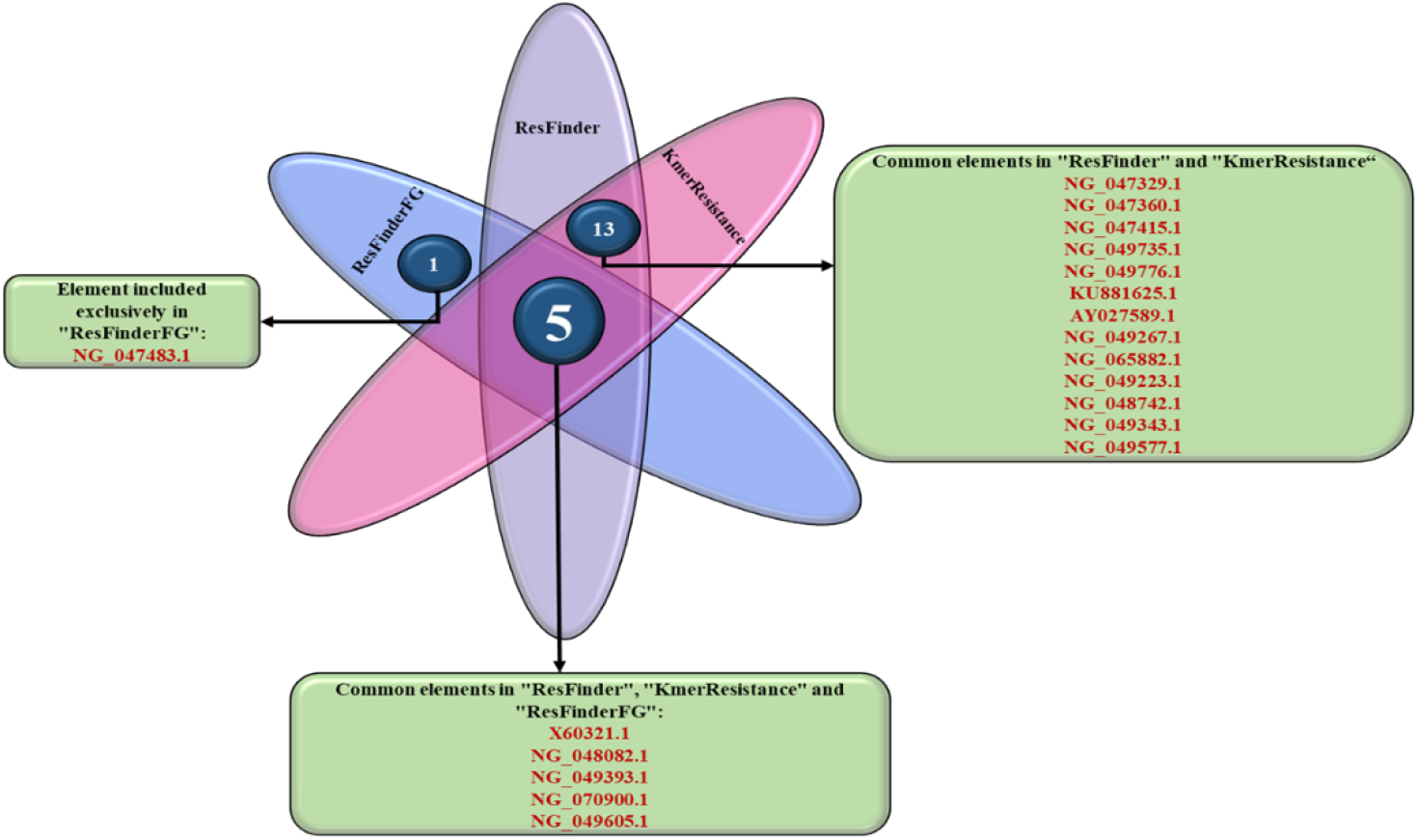
Venn diagram comparing the plasmid Accession Id results of ResFinder 4.1, KmerResistance2.2 and ResFinderFG 2.0

### Comparison based on the output Antibiotic Resistance Genes (ARGs)

KmerResistance detected 15 of the 16 AMR genes predicted by ResFinder, along with four additional resistance genes. KmerResistance did not predict one of the genes predicted by ResFinder (Figure 5). ResFinderFG, on the other hand, cannot detect any AMR genes and can only predict the AMR determinants.

**Figure 5.**
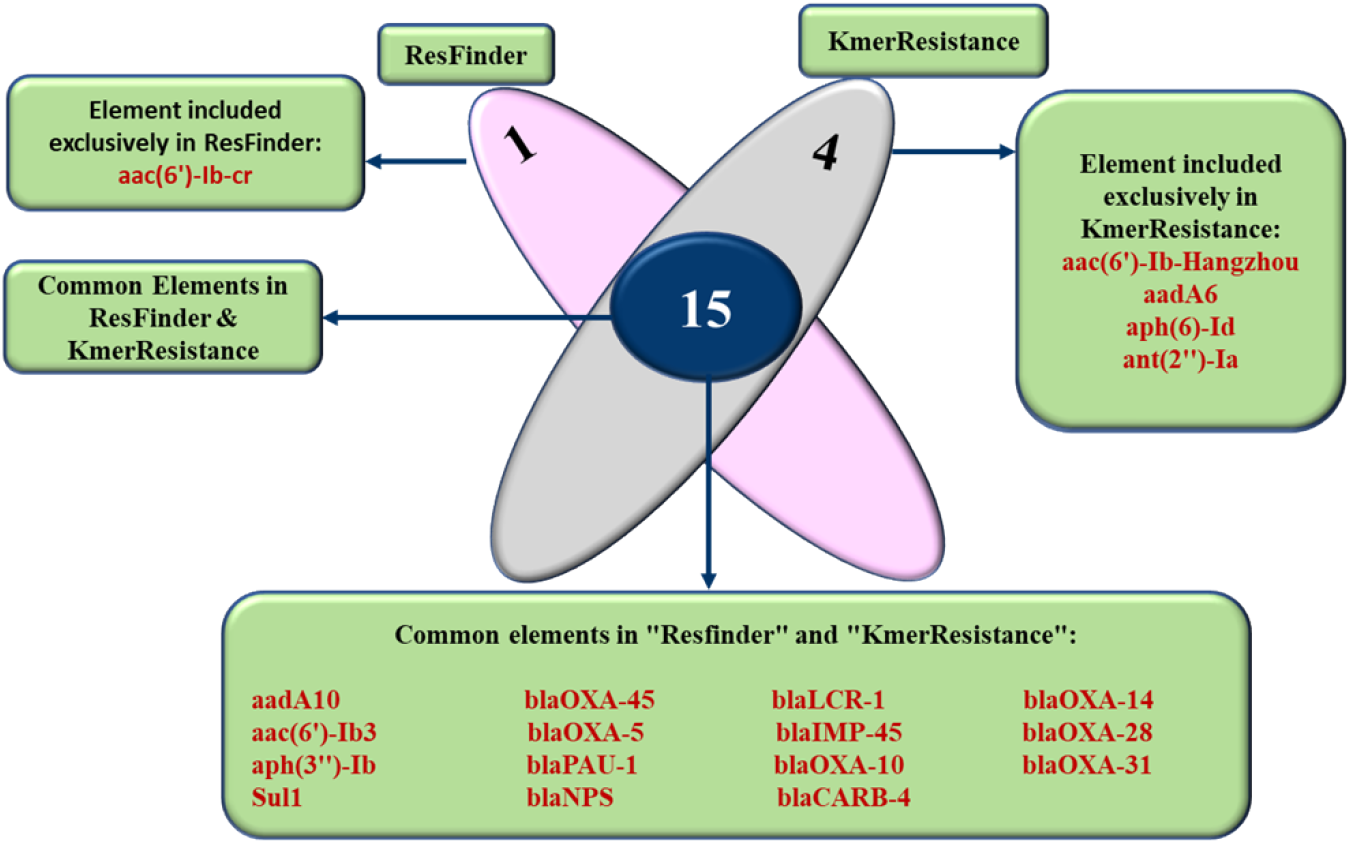
Venn diagram comparing the genes results of ResFinder 4.1 and KmerResistance 2.2

### Comparison based on the amount of information provided and result visualisation

The results of all three tools are made available through email. The outcomes of the analysis were displayed in a tabular manner in ResFinder, with the first table containing Antimicrobial, Class, WGS-predicted phenotype, and Genetic background. Other tables in ResFinder are organised by drug class, with each table containing various drug class information such as Resistance gene, Identity, Alignment Length/Gene Length, Position in reference, Contig or Depth, Position in contig, Phenotype, PMID, Accession no., and Notes. There is an extended output option at the bottom of the result page that displays the alignment result. The results for KmerResistance are provided in a single table which contains the following columns: Template, Score, Expected, Template Length, Q Value, P Value, Template Id, Template Coverage, Query Id, Query Coverage, Depth, and Depth Corr. The ResFinderFG result is displayed as a tabular box with numerous columns containing Hit name, Identity, Query/HSP, Contig, Position in contig, Drug treatment, and Accession no. Each of the three tools offers a variety of downloadable files containing various types of information (Table 3).

**Table 1.** Length and accession numbers of the plasmid sequences used.

| Sl no. | Accession no. | Plasmid length |
| --- | --- | --- |
| 1 | NZ_WTXS01000233.1 | 929 bps |
| 2 | NZ_WTXS01000231.1 | 988 bps |
| 3 | NZ_WTXR01000281.1 | 1,038 bps |
| 4 | NG_070900.1 | 1,001bps |
| 5 | NG_049735.1 | 995 bps |
| 6 | NG_049605.1 | 1,031 bps |
| 7 | NG_049577.1 | 1,001 bps |
| 8 | X60321.1 | 919 bps |
| 9 | D78374.1 | 1,014 bps |
| 10 | NZ_CP033773.1 | 1,089 bps |
| 11 | MN013162.1 | 1,022 bps |
| 12 | CP033773.1 | 1,089 bps |
| 13 | NG_065882.1 | 912 bps |
| 14 | KU881625.1 | 912 bps |
| 15 | AY027589.1 | 957 bps |
| 16 | NG_049776.1 | 983 bps |
| 17 | NG_049393.1 | 1,001 bps |
| 18 | NG_049343.1 | 928 bps |
| 19 | NG_049267.1 | 983 bps |
| 20 | NG_048742.1 | 1,067 bps |
| 21 | NG_048082.1 | 1,040 bps |
| 22 | NG_047483.1 | 1,082 bps |
| 23 | NG_047415.1 | 1,004 bps |
| 24 | NG_047360.1 | 969 bps |
| 25 | NG_047329.1 | 1,034 bps |
| 26 | NG_049223.1 | 938 bps |

**Table 2.** Details of tools for the prediction of drug resistance genes (2012-2022)

| Source | Description | Accessibility | Approach | Type | Year | Availability status | Ref. |
| --- | --- | --- | --- | --- | --- | --- | --- |
| **ARG-ANNOT** | It identifies current and potentially new antibiotic resistance (AR) genes in bacterial genomes. | Standalone | Assembly-based tools | Tool and database | 2014 | Archived last update in May 2018 | (20) |
| **ARGs-OAP** | An online analytic tool that uses an integrated structured ARG-database to discover antibiotic resistance genes in metagenomic data. | Web and/or standalone | Assembly-based tools | Tool | 2016 | Active | (21) |
| **ARIBA** | It is a tool for rapid genotyping antimicrobial resistance based on sequence reads. | Standalone | Assembly-based tools | Tool | 2017 | Active | (22) |
| **DeepArg** | A deep learning framework for identifying antibiotic resistance genes using metagenomic data. | Web | Read-based tools | Tool | 2018 | Active | (23) |
| **GROOT** | A tool for identifying Antibiotic Resistance Genes (ARGs) from metagenomic samples. | Standalone | Read-based tools | Tool | 2018 | Active | (24) |
| **KmerResistance** | KmerResistance links mapped genes with identified species of Whole Genome Sequencing (WGS) samples, allowing for gene identification in badly sequenced samples or highly accurate predictions for contaminated samples. | Web | Read-based tools | Tool | 2016 | Active | (25) |
| **NCBI-AMRFinder** | Using protein annotations and/or assembled nucleotide sequences, this tool can discover AMR genes, resistance-associated point mutations, and other kinds of genes. | Standalone | Assembly-based tools | Tool | 2018 | Active | (26) |
| **PATRIC** | It offers comprehensive data and analysis tools to help biomedical researchers study bacterial infectious illnesses. | Web | Read-based tools | Tool | 2016 | Active | (27) |
| **ResFinder** | It identifies acquired genes and/or chromosomal alterations mediating antimicrobial resistance in bacteria's complete or partial DNA sequence. | Web and standalone | Assembly-based and read based tool | Tool and database | 2012 | Active update regularly | (28) |
| **ResFinderFG** | Based on a functional metagenomic antibiotic resistance determinants database, it identifies a resistance phenotype. | Web | Assembly-based tool | Tool and database | 2016 | Active | Unpublished |
| **RGI** | It can be used to identify resistomes utilizing protein or nucleotide data using homology and SNP models. | Web and/or standalone | Assembly-based tools | Tool | 2015 | Active | (28) |
| **SEAR** | Create full-length, horizontally acquired Antibiotic Resistance Genes (ARGs) from sequencing data. It is intended for use in environmental metagenomics and microbiome research where the diversity and relative abundance of ARGs must be determined fast and conveniently. | Web and/or standalone (archived) | Read-based tools | Tool | 2015 | Archived | (29) |
| **ShortBRED** | ShortBRED is a high-specificity approach for profiling protein families of concern in shotgun meta-omic sequencing data. | Standalone | Read-based tools | Tool | 2015 | Active | (30) |
| **SRST2** | Rapid genomic monitoring for public health-care and clinical microbiology laboratories. | Standalone | Read-based tools | Tool | 2014 | Active | (31) |
| **SSTAR** | SSTAR offers rapid and accurate antimicrobial resistance (AR) surveillance using data from WGS. It can identify known AR genes as well as possible novel variations and shortened genes caused by internal stop codons. SSTAR can also detect changes and/or truncations in outer membrane porin gene sequences. | Standalone | Read-based tools | Tool | 2016 | Active | (32) |
| **ARGDIT** | It is an Antimicrobial Resistance Gene Database Validation and Integration Toolkit. | Standalone | Read-based tools | Tool | 2019 | Active | (33) |
| **LRE-Finder** | Finding the genes and mutations responsible for linezolid resistance | Web | Assembly-based tool | Tool and database | 2019 | Active | (34) |
| **IRIDA plugin AMR detection** | It is a pipeline that detects antimicrobial resistance genes using the RGI/CARD and starmr tools | Standalone | Assembly-based tool | Tool | 2019 | Active | (35) |
| **PARGT** | Software for predicting bacterial antimicrobial resistance | Standalone | Read-based tools | Tool | 2020 | Active | (36) |
| **HMD-ARG** | Annotating antibiotic resistance genes using hierarchical multi-task deep learning | Standalone | Read-based tools | Tool | 2021 | Active | (37) |
| **YunxiaoRen / ML-iAMR** | Antimicrobial resistance prediction using whole-genome sequencing and machine learning | Standalone | Read-based tools | Tool | 2022 | Active | (38) |

**Table 3.** AMR Tools and provided downloadable files.

| Tools | Output downloadable files |
| --- | --- |
| **ResFinder** | Phenotype table, Species specific phenotype table, Results as text (acquired AMR gene results), Resistance gene sequences, Hit in genome sequences, Results as tab separated file (acquired AMR gene results), Results as tab separated file (Chromosomal point mutation results) and Results as a text file (Chromosomal point mutation results) |
| **ResFinderFG** | Hit in genome sequences, Results as text, Resistance gene sequences and Results as tab separated file |
| **KmerResistance** | Resistance results, Species results, Full resistance results, Resistance alignment results, Resistance consensus results, Not-sam file and Log file |

### In detailed result of each tool

As a case study, we took 26 plasmid sequences and ran them through three different prediction tools to identify AMR factors.

### ResFinder 4.1

ResFinder 4.1 predicted that plasmid X60321.1 contains the majority of AMR genes. It carries two AMR genes, aac(6’)-Ib3 resistance to aminoglycosides and aac(6’)-Ib-cr resistance to fluoroquinolone classes of antibiotics. The presence of AMR genes was found in 18 of the 25 plasmid sequences tested in this study, with no resistance gene found for thirteen of the drug classes: Colistin, Disinfectant, Fosfomycin, Fusidic acid, Glycopeptide, MLS (Macrolide, Lincosamide, and Streptogramines), Nitroimidazole, Oxazolidinone, Phenicol, Pseudomonic acid, Rifampicin, Tetracycline, and Trimethoprim. A fair number of AMR gene homologues were identified in the other four drug classes (Supplementary Table 1). The drug classes in which AMR genes were detected were beta-lactams (13 out of 19; 68.42%), followed by aminoglycosides (21.05%), fluoroquinolones (5.2%), and sulphonamides (5.2%). There were 16 number of different acquired AMR genes found to be resistant to the four drug classes, with aminoglycosides and beta-lactams showing the highest frequency (Table 4). The most common genes are aadA10 (2/19), blaNPS (2/19), and blaPAU-1 (2/19), each of which was found to be present in 10.52% of the outcomes observed (Figure. 6)

**Table 4.**
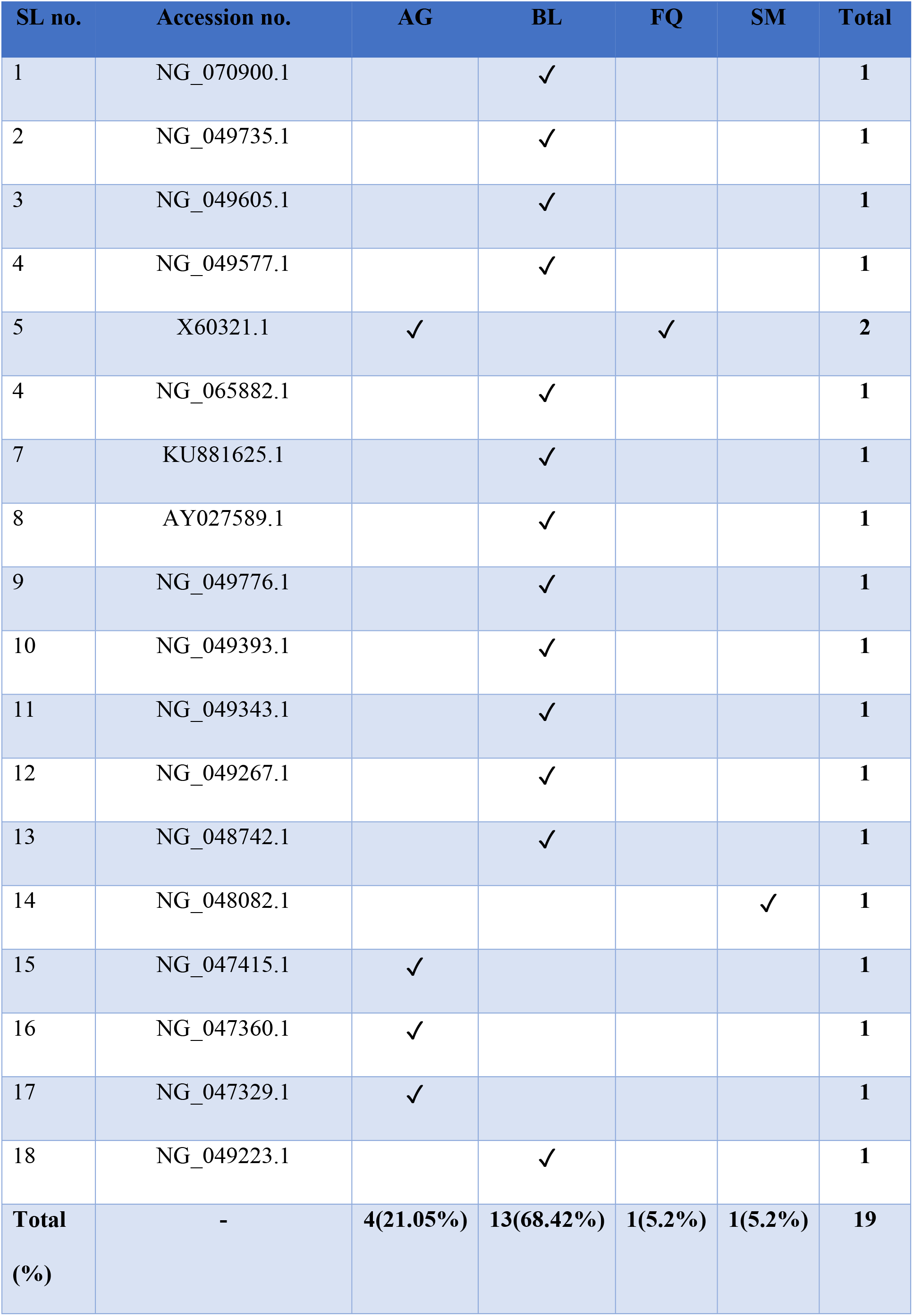
Resistance to specific antimicrobial drug classes was identified in the input plasmid sequences. The presence of various numbers of AMR genes to four drug classes was observed in the 18 plasmids listed. (AG = aminoglycoside; BL = beta-lactam; FQ = fluoroquinolone; SM = sulphonamide)

**Figure 6.**
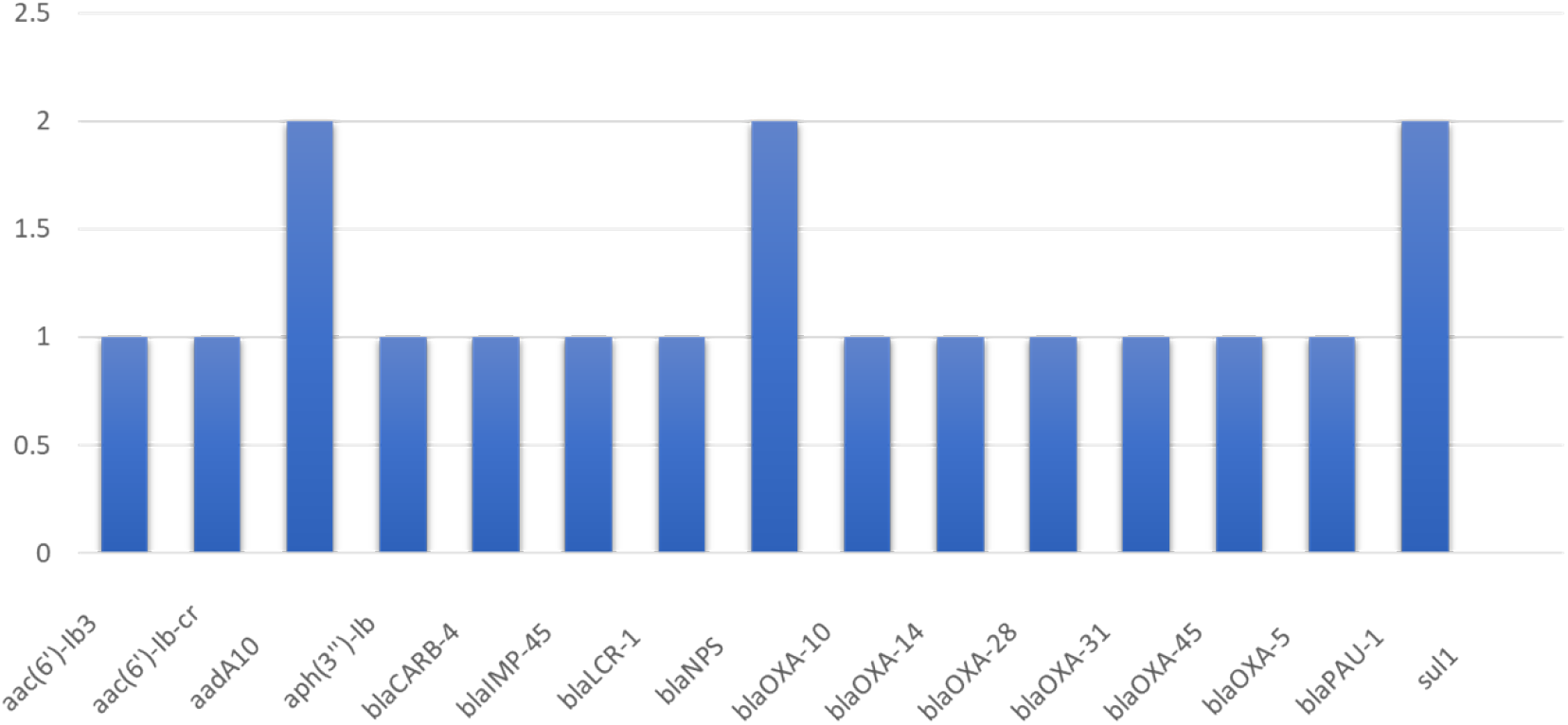
ResFinder 4.1 revealed the prevalence of 16 AMR genes found in 18 Pseudomonas aeruginosa plasmids.

### KmerResistance 2.2

For an in-silico analysis of AMR genes, the sequences were uploaded to the KmerResistance database. Eight plasmid template genes were linked to other Gram-negative bacteria, while one plasmid template gene was linked to a Gram-positive bacteria, indicating that they could be the source of acquisition (Supplementary Table 2). Two Escherichia coli, one each of Klebsiella pneumoniae, Comamonas testosterone, Acinetobacter sp., Pseudomonas putida, Achromobacter xylosoxidans, and Providencia sp. linked with Gram-negative strain. Glutamicibacter nicotianae Gram-positive strain is associated with one of the plasmids. Out of 25 input plasmid sequences, it predicted 19 ARGs in 13 plasmids.

### ResFinderFG 2.0

In ResFinderFG 2.0 three distinct ARD families were identified in six input plasmid sequences. The most common was beta-lactamase, which was found in four different input plasmid sequences. Three of the four plasmids with beta-lactamase ARDs conferred resistance to ampicillin and one conferred resistance to piperacillin. Others include one dihydropteroate synthase (dpr) and one aminoglycoside acetyl-transferase (AAC), as both are resistant to Sulfamethazine and Amikacin respectively (Supplementary Table 3).

## Discussion

Currently, the use of sequencing technology is revolutionizing practically every component of biological study. In the field of infectious diseases, scientific discoveries, as well as diagnostic and outbreak investigations, are developing at a rapid pace. The ability to interpret sequencing data and the benefit of quick development, on the other hand, is not evenly distributed between institutions and countries (39,40). We choose CGE tools because its goal is to give access to bioinformatics tools for those with limited knowledge, allowing all countries, institutions, and individuals to benefit from new sequencing technology. It is believed that by doing so, it will encourage more open data sharing around the world and give equal advantages to all. CGE is fully non-profitable and provides a variety of free online bioinformatics services (19).

The AMR prediction methods have been built using DNA or amino acid sequence data. The presence or absence of software for searching within an AMR determinant database, which can be precise to a tool or replicated from other resources, the type of input data accepted, and the search approach used, which can be alignment or mapping, are all factors that distinguish bioinformatics resources. Each tool has its own set of capabilities and limitations when it comes to AMR prediction, which includes the identification of AMR sequences, ARGs, the volume of information produced, and visualisation. As the web-based application is one that uses a website as its interface or front-end. It has the potential to provide competitive benefits over traditional software-based systems by providing researchers to streamline data and information at a lower cost, time, and maintenance (19). Using a regular browser, users can quickly access the application across any computer with internet access. Functionality and features were the two main elements that were prioritized. From this study, a non-bioinformatician or non-technician can gain subject-specific knowledge as well as determine which tool is appropriate for their specific work. The table below compares the three tools used for identifying drug resistance genes and resistance plasmids as well as the amount of information they provide and the way their results are displayed. (Table 5).

**Table 5.**
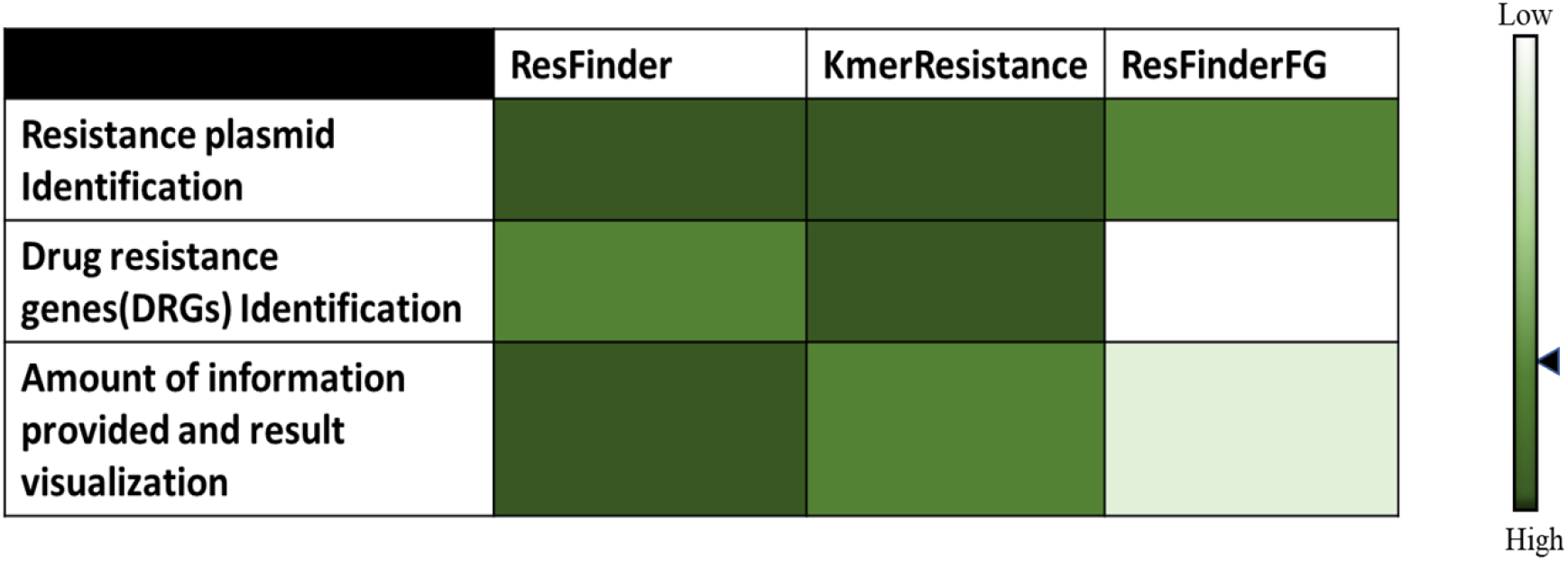
Empirical evaluation of three different AMR prediction tools.

### The description and significant observations of each selected web-based tools have been discussed below

ResFinder is a web-based tool that finds chromosomal alterations that promote antibiotic resistance in bacteria’s whole or partial DNA sequence and identifies acquired antimicrobial resistance genes in the whole-genome data using BLAST. As input, this tool accepts pre-assembled, whole or partial genomes, along with fragmented sequence reads from four distinct sequencing technologies. It is accessible at (https://cge.food.dtu.dk/services/ResFinder/). It is also constantly being updated whenever different resistance genes are discovered (41). Here, we found that the database showed a restriction that stated it only looked for acquired genes and did not detect chromosomal mutations. Since new resistance genes are continuously discovered, it may be necessary to confirm the presence or absence of identified AMR genes phenotypically. According to a study (42), genotyping using aligned whole-genome sequences is a practical substitute for surveillance based on phenotypic antimicrobial susceptibility due to the high concordance (99.74%) between phenotypic and predicted whole-genome sequence antimicrobial susceptibility. One of the three most common genes, the aadA10 gene, was found to be resistant to the drug class aminoglycoside, while the other two genes, blaNPS and blaPAU-1, were found to be resistant to the beta-lactam drug class. AadA10 is a class 1 integron containing gene cassette, which suggests that rather than transposition, the resistance determinants from one plasmid to another plasmid were moved by recombination between two class 1 integrons (43). The ability of integrons in bacteria to acquire new cassettes and recombine cassette rows emphasises the adaptability of integron diversity. It is necessary to be aware of what other integron-mediated traits, such as increased resistance to antimicrobials, virulence, or pathogenicity, might affect human health in the future given their capacity to quickly spread resistance phenotypes. There is an urgent need for control integrons and cassette formation (44). Tauch et al., 2003, found that blaNPS can rapidly transfer from one species to another (45). According to Subedi et al., 2018, environmental resistance gene pools contain blaNPS, which can be acquired and maintained in clinical isolates. (46). In a prior study, it was discovered that a clinical isolate of *Pseudomonas aeruginosa* contained a transferable plasmid containing the gene known as blaPAU-1, which is connected to the mobile genetic element (47). Considering the high ubiquity of beta-lactam resistance genes, it is preferable to monitor the level of antibiotic resistance and resistance genes in patients with *Pseudomonas aeruginosa* infection (48). In this study, we found that *Pseudomonas aeruginosa* plasmids contain many beta-lactam resistance genes, a number of which are found in a single plasmid. This leads us to the conclusion that beta-lactam resistance genes are rapidly spreading through plasmids, and the need of the hour is to control their spread.

KmerResistance (https://cge.food.dtu.dk/services/KmerResistance/) is primarily based on KmerFinder, which has been developed for typing of bacteria with raw WGS data. KmerResistance and KmerFinder search for co-occurring k-mers between such a query genome and a resistance gene database. KmerResistance, like KmerFinder, biases the threshold based on the quality of the data, as shown by the coverage as well as the depth of the detected species genome. Since this k-mers in this scenario are dispersed across the total sample, we can estimate both depth and coverage. Unlike KmerFinder, KmerResistance may generate an outcome for species prediction in addition to getting acquired antimicrobial resistance genes.(49). This database improves on poor-quality assembly by using k-mers to map raw whole-genome sequence data against reference databases and species. (Fragments of a DNA sequence of length k) (28). Additionally, it can find host or template genes. The KmerResistance database displays the resistance genes but not the drug classes as an analysis output, even though being claimed to be more precise than ResFinder. As a result, comparisons were restricted to resistance genes that were present in both databases rather than an overall assessment of their sensitivity. ResFinder and ResFinderFG accept input sequences in a single input file and provide results based on each sequence, while KmerResistance requires us to provide the sequence file separately and perform separate executions. If we provide the input sequence to KmerResistance in a single file, the results are very perplexing and unrelated to the input sequence.

The ResFinderFG (https://cge.food.dtu.dk/services/ResFinderFG/) approach is based on databases containing sequences detected by functional metagenomics but not represented in existing databases constructed mostly from antibiotic-resistant genes in clinical isolates. It identifies a resistant phenotype in general (50). One of the main causes of beta-lactam resistance in *Pseudomonas aeruginosa* is the increased prevalence of beta-lactamase (48). Beta-lactamase enzymes render beta-lactam antibiotics ineffective by hydrolyzing the peptide bond of the characteristic four-membered beta-lactam ring. The bacterium gains resistance after the antibiotic is rendered inactive. Over 300 beta-lactamase enzymes have been described so far, with numerous kinetic, structural, computational, and mutagenesis studies. The threat posed by more and more powerful beta-lactamases to antimicrobial therapy is only going to increase (51). The scientific and medical communities are only one step ahead and must continue to put forth diligent effort to prevent being overcome by the difficult and rapidly rising nosocomial pathogen resistance.

## Conclusion

A variety of freely accessible tools are available for the prediction of AMR determinants. A Comparison of these tools will aid in the expansion of global pathogen monitoring and AMR tracking based on genomic information. Since AMR tools are used to accomplish various tasks, it would be unfair and difficult to compare them directly. We provided a case study using bioinformatics tools that allow researchers to predict AMR. Here we discovered that ResFinder is very advantageous for the prediction of resistant plasmid identification and provides a large amount of information with better visualisation, whereas KmerResistance is quite good for resistance plasmid identification, information regarding related species and the template gene, as well as predicting ARGs. ResFinderFG does not provide any information about ARGs and provides relatively lesser information than the other two tools, although the visualisation is superior to KmerResistance. Each has its own unique strengths and weaknesses. Our research provides a thorough overview of the results obtained by three different tools, allowing users to select the best tool for their needs.

## Materials and methods

The framework employed in this study is described in the subsequent sections, and the comprehensive methodology is depicted in the figure below (Figure 2).

### Validation dataset

*Pseudomonas aeruginosa* is a pathogenic Gram-negative bacterium that is becoming more common in infections caused by Multidrug-resistant (MDR) and extensively drug-resistant (XDR) strains, limiting available effective treatments. Plasmids contribute significantly in antibiotic resistance because they are the means by which resistance genes are captured and then disseminated. *Pseudomonas aeruginosa* plasmid nucleotide sequences were acquired using the key phrase *“Pseudomonas aeruginosa”* from the nucleotide database of the NCBI (https://www.ncbi.nlm.nih.gov/nuccore/). The organism was then chosen as *Pseudomonas aeruginosa*, the species as bacteria, the molecular type as genomic DNA/RNA, the sequence type as nucleotide, genetic compartment as plasmid, and the length from 900bps to 1100bps.

In order to obtain a standard range, the length of the sequence was taken into account by the Romaniuk et al., 2019 article (18). We obtained 26 *Pseudomonas aeruginosa* plasmid sequences from the aforementioned screening, which are further considered for the AMR prediction. The table (Table 1) below includes NCBI accession numbers, plasmid length and species of the retrieved sequences.

### Segregation of in silico AMR determination tools

The development of online databases and bioinformatics tools has been required for drug-resistance gene prediction. After conducting a scientific literature search from 2012 to 2022, Forty-seven freely accessible bioinformatics tools for identifying AMR determinants were discovered, including the most commonly used tools listed below (Table 2).

Center for Genomic Epidemiology (CGE) is fully non-profitable and provides a variety of free online bioinformatics services. The Technical University of Denmark (DTU) provides core funding, as well as funding from a variety of public and commercial sources (19). The Center for Genomic Epidemiology offers 38 services, including nine phenotyping tools: ResFinder, ResFinderFG, LRE-finder, KmerResistance, PathogenFinder, VirulenceFinder, Restriction-ModificationFinder, SPIFinder, and ToxFinder (Figure 3). From the mentioned nine phenotypic tools, we selected three tools with versions (ResFinder 4.1, KmerResistance 2.2, and ResFinderFG 2.0) based on similarities of their objective i.e., to find out the resistance factors. Here we did not include LRE-Finder in our study because the input data format is FASTQ sequence, it only predicts AMR for a single species i.e, Enterococcus faecalis, and only identifies acquired linezolid resistance genes without considering other antibiotics.

### Steps and parameters setup for segregated tools

ResFinder 4.1 searched the database for all seventeen classes of antibiotic drugs, regardless of the target region. The sequences were entered into the database, and the acquired resistance gene testing parameters were adjusted to predict resistance genes for all seventeen drug classes provided by the server. The minimum percentage identity was set to 90%, with perfect alignment set to 100%. The percentage of identity was computed by counting the number of identical nucleotides between the best-matching resistance gene in the database and the equivalent sequence in the plasmid. The tool was run with the aforementioned parameters, and the results were recorded.

The scoring method in KmerResistance 2.2 was species identification on maximum query coverage, and also the host database was set to the bacterial plasmid. The gene database was set to resistant genes, and the identity threshold was left at 70%, with a depth correction threshold of 10%. The AMR genes identified in the resulting output were recorded and the host organism and template sequence were also noted.

The functional genomics database ResFinderFG 2.0 uses functional metagenomic antibiotic resistance determinants to identify resistance phenotypes. The percentage identity setting was set to 98%, along with the minimum query length was set to 60%. The read type used was ‘assembled contigs/genomes,’ and the sequences were screened for all the thirteen antibiotic resistance determinant (ARD) families present in the database.

## Acknowledgement

The author acknowledges and thank the Center for Genomic Epidemiology in Denmark for providing web tools.

## References

1. Aslam B, Wang W, Arshad MI, Khurshid M, Muzammil S, Rasool MH, et al. Antibiotic resistance: a rundown of a global crisis. Vol. 11, Infection and Drug Resistance. 2018.

2. Pagès JM, Amaral L. Mechanisms of drug efflux and strategies to combat them: Challenging the efflux pump of Gram-negative bacteria. Vol. 1794, Biochimica et Biophysica Acta - Proteins and Proteomics. 2009.

3. Hutchings M, Truman A, Wilkinson B. Antibiotics: past, present and future. Vol. 51, Current Opinion in Microbiology. 2019.

4. Ferri M, Ranucci E, Romagnoli P, Giaccone V. Antimicrobial resistance: A global emerging threat to public health systems. Crit Rev Food Sci Nutr. 2017;57(13).

5. Jasovský D, Littmann J, Zorzet A, Cars O. Antimicrobial resistance—a threat to the world’s sustainable development. Vol. 121, Upsala Journal of Medical Sciences. 2016.

6. Zhang AN, Gaston JM, Dai CL, Zhao S, Poyet M, Groussin M, et al. An omics-based framework for assessing the health risk of antimicrobial resistance genes. Nat Commun. 2021;12(1).

7. Lerminiaux NA, Cameron ADS. Horizontal transfer of antibiotic resistance genes in clinical environments. Can J Microbiol. 2019;65(1).

8. Zankari E, Hasman H, Cosentino S, Vestergaard M, Rasmussen S, Lund O, et al. Identification of acquired antimicrobial resistance genes. Journal of Antimicrobial Chemotherapy. 2012;67(11).

9. Gajic I, Kabic J, Kekic D, Jovicevic M, Milenkovic M, Mitic Culafic D, et al. Antimicrobial Susceptibility Testing: A Comprehensive Review of Currently Used Methods. Vol. 11, Antibiotics. 2022.

10. Motro Y, Moran-Gilad J. Next-generation sequencing applications in clinical bacteriology. Vol. 14, Biomolecular Detection and Quantification. 2017.

11. Nayak DSK, Mahapatra S, Swarnkar T. Gene selection and enrichment for microarray data—A comparative network based approach. In: Advances in Intelligent Systems and Computing. 2018.

12. Khan ZA, Siddiqui MF, Park S. Current and emerging methods of antibiotic susceptibility testing. Vol. 9, Diagnostics. 2019.

13. Mahfouz N, Ferreira I, Beisken S, von Haeseler A, Posch AE. Large-scale assessment of antimicrobial resistance marker databases for genetic phenotype prediction: A systematic review. Vol. 75, Journal of Antimicrobial Chemotherapy. 2020.

14. Hendriksen RS, Bortolaia V, Tate H, Tyson GH, Aarestrup FM, McDermott PF. Using Genomics to Track Global Antimicrobial Resistance. Vol. 7, Frontiers in Public Health. 2019.

15. Seoane A, Bou G. Bioinformatics approaches to the study of antimicrobial resistance. Revista Espanola de Quimioterapia. 2021;34.

16. Hendriksen RS, Bortolaia V, Tate H, Tyson GH, Aarestrup FM, McDermott PF. Using Genomics to Track Global Antimicrobial Resistance. Vol. 7, Frontiers in Public Health. 2019.

17. Köser CU, Ellington MJ, Peacock SJ. Whole-genome sequencing to control antimicrobial resistance. Vol. 30, Trends in Genetics. 2014.

18. Romaniuk K, Styczynski M, Decewicz P, Buraczewska O, Uhrynowski W, Fondi M, et al. Diversity and horizontal transfer of antarctic pseudomonas spp. plasmids. Genes (Basel). 2019;10(11).

19. Thomsen MCF, Ahrenfeldt J, Cisneros JLB, Jurtz V, Larsen MV, Hasman H, et al. A bacterial analysis platform: An integrated system for analysing bacterial whole genome sequencing data for clinical diagnostics and surveillance. PLoS One. 2016;11(6).

20. Gupta SK, Padmanabhan BR, Diene SM, Lopez-Rojas R, Kempf M, Landraud L, et al. ARG-annot, a new bioinformatic tool to discover antibiotic resistance genes in bacterial genomes. Antimicrob Agents Chemother. 2014;58(1).

21. Yang Y, Jiang X, Chai B, Ma L, Li B, Zhang A, et al. ARGs-OAP: Online analysis pipeline for antibiotic resistance genes detection from metagenomic data using an integrated structured ARG-database. Bioinformatics. 2016;32(15).

22. Hunt M, Mather AE, Sánchez-Busó L, Page AJ, Parkhill J, Keane JA, et al. ARIBA: Rapid antimicrobial resistance genotyping directly from sequencing reads. Microb Genom. 2017;3(10).

23. Arango-Argoty G, Garner E, Pruden A, Heath LS, Vikesland P, Zhang L. DeepARG: A deep learning approach for predicting antibiotic resistance genes from metagenomic data. Microbiome. 2018;6(1).

24. Rowe WPM, Winn MD. Indexed variation graphs for efficient and accurate resistome profiling. Bioinformatics. 2018;34(21).

25. Clausen PTLC, Zankari E, Aarestrup FM, Lund O. Benchmarking of methods for identification of antimicrobial resistance genes in bacterial whole genome data. Journal of Antimicrobial Chemotherapy. 2016;71(9).

26. Feldgarden M, Brover V, Haft DH, Prasad AB, Slotta DJ, Tolstoy I, et al. Using the NCBI AMRFinder Tool to Determine Antimicrobial Resistance Genotype-Phenotype Correlations Within a Collection of NARMS Isolates. bioRxiv. 2019.

27. Wattam AR, Davis JJ, Assaf R, Boisvert S, Brettin T, Bun C, et al. Improvements to PATRIC, the all-bacterial bioinformatics database and analysis resource center. Nucleic Acids Res. 2017;45(D1).

28. Zankari E, Hasman H, Cosentino S, Vestergaard M, Rasmussen S, Lund O, et al. Identification of acquired antimicrobial resistance genes. Journal of Antimicrobial Chemotherapy. 2012;67(11).

29. Rowe W, Baker KS, Verner-Jeffreys D, Baker-Austin C, Ryan JJ, Maskell D, et al. Search engine for antimicrobial resistance: A cloud compatible pipeline and web interface for rapidly detecting antimicrobial resistance genes directly from sequence data. PLoS One. 2015;10(7).

30. Kaminski J, Gibson MK, Franzosa EA, Segata N, Dantas G, Huttenhower C. High-Specificity Targeted Functional Profiling in Microbial Communities with ShortBRED. PLoS Comput Biol. 2015;11(12).

31. Inouye M, Dashnow H, Raven LA, Schultz MB, Pope BJ, Tomita T, et al. SRST2: Rapid genomic surveillance for public health and hospital microbiology labs. Genome Med. 2014;6(11).

32. de Man TJB, Limbago BM. SSTAR, a Stand-Alone Easy-To-Use Antimicrobial Resistance Gene Predictor. mSphere. 2016;1(1).

33. Chiu JKH, Ong RTH. ARGDIT: A validation and integration toolkit for Antimicrobial Resistance Gene Databases. Bioinformatics. 2019;35(14).

34. Hasman H, Clausen PTLC, Kaya H, Hansen F, Knudsen JD, Wang M, et al. LRE-Finder, a Web tool for detection of the 23S rRNA mutations and the optrA, cfr, cfr(B) and poxtA genes encoding linezolid resistance in enterococci from whole-genome sequences. Journal of Antimicrobial Chemotherapy. 2019;74(6).

35. Matthews TC, Bristow FR, Griffiths EJ, Petkau A, Adam J, Dooley D, et al. The Integrated Rapid Infectious Disease Analysis (IRIDA) Platform. bioRxiv. 2018;

36. Chowdhury AS, Call DR, Broschat SL. PARGT: a software tool for predicting antimicrobial resistance in bacteria. Sci Rep. 2020;10(1).

37. Li Y, Xu Z, Han W, Cao H, Umarov R, Yan A, et al. HMD-ARG: hierarchical multi-task deep learning for annotating antibiotic resistance genes. Microbiome. 2021;9(1).

38. Ren Y, Chakraborty T, Doijad S, Falgenhauer L, Falgenhauer J, Goesmann A, et al. Prediction of antimicrobial resistance based on whole-genome sequencing and machine learning. Bioinformatics. 2022;38(2).

39. Gardy JL, Loman NJ. Towards a genomics-informed, real-time, global pathogen surveillance system. Vol. 19, Nature Reviews Genetics. 2018.

40. Gilchrist CA, Turner SD, Riley MF, Petri WA, Hewlett EL. Whole-genome sequencing in outbreak analysis. Clin Microbiol Rev. 2015;28(3).

41. Bortolaia V, Kaas RS, Ruppe E, Roberts MC, Schwarz S, Cattoir V, et al. ResFinder 4.0 for predictions of phenotypes from genotypes. Journal of Antimicrobial Chemotherapy. 2020;75(12).

42. Zankari E, Hasman H, Kaas RS, Seyfarth AM, Agersø Y, Lund O, et al. Genotyping using whole-genome sequencing is a realistic alternative to surveillance based on phenotypic antimicrobial susceptibility testing. Journal of Antimicrobial Chemotherapy. 2013;68(4).

43. Partridge SR, Collis CM, Hall RM. Class 1 integron containing a new gene cassette, aadA10, associated with Tn1404 from R151. Antimicrob Agents Chemother. 2002;46(8).

44. Sabbagh P, Rajabnia M, Maali A, Ferdosi-Shahandashti E. Integron and its role in antimicrobial resistance: A literature review on some bacterial pathogens. Vol. 24, Iranian Journal of Basic Medical Sciences. 2021.

45. Tauch A, Schlüter A, Bischoff N, Goesmann A, Meyer F, Pühler A. The 79,370-bp conjugative plasmid pB4 consists of an IncP-1β backbone loaded with a chromate resistance transposon, the strA-strB streptomycin resistance gene pair, the oxacillinase gene blaNPS-1, and a tripartite antibiotic efflux system of the resistance-nodulation-division family. Molecular Genetics and Genomics. 2003;268(5).

46. Subedi D, Vijay AK, Kohli GS, Rice SA, Willcox M. Nucleotide sequence analysis of NPS-1 β-lactamase and a novel integron (In1427)-carrying transposon in an MDR Pseudomonas aeruginosa keratitis strain. Vol. 73, Journal of Antimicrobial Chemotherapy. 2018.

47. Wang J, Xu T, Ying J, Zhou W, Chen Q, Qian C, et al. PAU-1, a novel plasmid-encoded ambler class a β-lactamase identified in a clinical Pseudomonas aeruginosa isolate. Infect Drug Resist. 2019;12.

48. Haghighi S, Goli HR. High prevalence of blaVEB, blaGES and blaPER genes in beta-lactam resistant clinical isolates of Pseudomonas aeruginosa. AIMS Microbiol. 2022;8(2):153–66.

49. Clausen PTLC, Zankari E, Aarestrup FM, Lund O. Benchmarking of methods for identification of antimicrobial resistance genes in bacterial whole genome data. Journal of Antimicrobial Chemotherapy. 2016;71(9).

50. Blake KS, Choi JH, Dantas G. Approaches for characterizing and tracking hospital-associated multidrug-resistant bacteria. Vol. 78, Cellular and Molecular Life Sciences. 2021.

51. Majiduddin FK, Materon IC, Palzkill TG. Molecular analysis of beta-lactamase structure and function. International Journal of Medical Microbiology. 2002;292(2).

